# Dissecting the interactions of PINK1 with the TOM complex in depolarized mitochondria

**DOI:** 10.1101/2022.01.13.476189

**Authors:** Klaudia K. Maruszczak, Martin Jung, Shafqat Rasool, Jean-François Trempe, Doron Rapaport

## Abstract

Mitochondria dysfunction is involved in the pathomechanism of many illnesses including Parkinson’s disease. PINK1, which is mutated in some cases of familiar Parkinsonism, is a key component in the degradation of damaged mitochondria by mitophagy. The accumulation of PINK1 on the mitochondrial outer membrane (MOM) of compromised organelles is crucial for the induction of mitophagy, but the molecular mechanism of this process is still unresolved. Here, we investigate the association of PINK1 with the TOM complex. We demonstrate that PINK1 heavily relies on the import receptor TOM70 for its association with mitochondria and directly interacts with this receptor. The structural protein TOM7 appears to play only a moderate role in PINK1 association with the TOM complex, probably due to its role in stabilizing this complex. PINK1 requires the TOM40 pore lumen for its stable interaction with the TOM complex and apparently remains there during its further association with the MOM. Overall, this study provides new insights on the role of the individual TOM subunits in the association of PINK1 with the MOM of depolarized mitochondria.

## Introduction

Mitochondria are versatile organelles that form tubular networks within eukaryotic cells. They are highly dynamic structures undergoing fission and fusion processes to adjust to existing conditions and to maintain cells in a healthy state (Ni *et al.*, 2015). Mitochondria also harbour many metabolic pathways and have a crucial role in supplying cells with energy in the form of ATP (Bertram *et al.*, 2006). Being critical for cellular homeostasis, mitochondria have to be continuously monitored by different quality control proteins and mechanisms. One of these mechanisms relies on mitochondrial autophagy (mitophagy) that is mediated by the proteins PINK1 and Parkin (Vincow *et al.*, 2013). In fully functional mitochondria, PINK1 is imported into the organelle where the inner membrane protease, PARL, cleaves it within the transmembrane domain (Greene *et al.*, 2012). Such processed PINK1 is subsequently retro-translocated into the cytosol and degraded by the proteasome (Yamano & Youle, 2013). In contrast, upon mitochondrial depolarization, PINK1 is accumulated at the mitochondrial outer membrane (MOM), where it phosphorylates itself and ubiquitin moieties conjugated to MOM proteins. Additionally, it phosphorylates also Parkin (E3 Ub ligase), which in turn ligates additional ubiquitin molecule to the previously phosphorylated ubiquitin (Narendra *et al.*, 2010; Matsuda *et al.*, 2010). This process leads to the formation of poly-ubiquitin chains and generation of positive feedback loop that eventually result in the specific elimination of compromised mitochondria by mitophagy. Hence, the association of PINK1 with the MOM is crucial for the whole mitophagy process. Dysfunctional mitophagy, and specifically mutations in PINK1 and/or Parkin, can lead to the development of neurodegenerative diseases with Parkinson’s disease as the main example (Morais *et al.*, 2009).

So far, the import of PINK1 into polarized mitochondria has been elucidated in some detail (Hasson et al., 2013; Nguyen et al., 2016; Yamano and Joule, 2013). In contrast, much less is known about its recognition and the subsequent integration into the MOM in depolarized organelles. Previous studies reported on the importance of the translocase of the mitochondrial outer membrane (TOM complex) for the integration of PINK1 into the MOM. For example, Lazarou et al. (2012) showed that PINK1 accumulates in the MOM in the form of high molecular weight complexes with the TOM complex. Later, Okatsu et al. (2013) discovered that PINK1 forms dimers in such complexes and is found there in its phosphorylated form. Considering these findings, a model was suggested in which the TOM complex facilitates the accurate orientation of the dimeric PINK1 so that intermolecular phosphorylation and subsequent activation can occur (Okatsu *et al.*, 2013). The structure of the *Tribolium castaneum* (Tc) PINK1 cytosolic domain revealed how dimerization enables *trans* autophosphorylation at Ser228 and suggests that anchoring on the TOM complex via an N-terminal helix is critical for PINK1 activation (Rasool et al. 2021).

Additional studies investigate the specific contribution of individual components of the TOM complex. For example, two studies proposed the relevance of the structural subunit TOM7, as being a “side gate” for PINK1 membrane insertion (Hasson *et al.*, 2013, Sekine *et al.*, 2019). Accordingly, addition of an uncoupler to HeLa cells lacking TOM7 failed to induce PINK1 accumulation at the MOM (Sekine *et al.*, 2019). In addition, both receptors of the TOM complex, TOM20 and TOM70 were suggested by two different studies to play an important role in PINK1 recognition at the MOM. Zhang et al. (2019) reported a drastic effect of TOM20 inhibition by celastrol on PINK1 association with mitochondria, and Kato et al. (2013) showed that PINK1 relies on the TOM70 receptor for its import into healthy mitochondria. Collectively, it seems that various subunits of the TOM complex contribute by an undefined way to the association of PINK1 with the MOM.

In addition to such trans elements, cis sequences within PINK1 itself were found to be important for its integration into the MOM. The outer mitochondrial membrane localization signal (OMS), which comprises a weak hydrophobic segment localized N-terminally to the transmembrane domain of PINK1 and encompasses amino acids residues 70-95 was reported to mediate association with the MOM (Okatsu *et al.*, 2015). Along this line, Sekine et al. (2019) have proposed that both TOM7 and OMS are crucial for PINK1 retention, with TOM7 mediating the lateral release of the protein from the TOM40 channel. Despite this progress, the location of PINK1 at the MOM of depolarized mitochondria and the factors that contribute to this positioning are only partially resolved.

Here, we investigated the initial import steps of PINK1 into depolarized mitochondria. We demonstrate that PINK1 heavily relies on TOM70 for its assembly into depolarized organelles and directly interacts with this receptor. Our findings further suggest that the accessory protein, TOM7, plays only a moderate role in PINK1 association with the TOM complex, probably due to its contribution to the stability of this complex. Importantly, PINK1 requires the TOM40 pore lumen for its association with the TOM complex and apparently remains there during its association with the MOM. Overall, this study provides new insights on the role of the TOM complex in the association of PINK1 with the MOM of depolarized mitochondria.

## Results

### PINK1 associates with the TOM complex upon depolarization of mitochondria

PINK1 accumulates at the MOM upon deprivation of mitochondrial inner membrane potential (ΔΨm). As part of our effort to dissect this process, we aimed to use a specific assay to monitor the association of PINK1 with the MOM. It was previously reported that PINK1 associates with the TOM complex upon CCCP treatment and forms a stable 700 kDa species that can be analysed by blue native (BN)-PAGE (Lazarou *et al.*, 2012). Hence, we decided to use this association as a readout for the productive association of PINK1 with the TOM complex. To validate that the detected 700 kDa band represents indeed an adduct of PINK1 and the TOM complex, we performed *in organello* import assay with radiolabelled PINK1 and mitochondria isolated from HeLa cells. As anticipated, when the import reactions were solubilized with digitonin and analysed by BN-PAGE, we observed a band at ca. 700 kDa only if mitochondria were depolarized before (Figure 1a). To verify the identity of this band, we aimed to immunodeplete the TOM complex from the digitonin suspension by addition of an antibody against TOM22 conjugated to protein A beads. This procedure allowed us to deplete the TOM complex, as analysed by BN-PAGE, from the organelle’s lysate (Figure 1b). In agreement with previous studies (Lazarou *et al.*, 2012; Hasson et al., 2013), this treatment removed also the 700 kDA from the lysate and in parallel resulted in binding of the PINK1-TOM adduct to the anti-TOM22 beads (Figure 1a). Accordingly, we detected TOM40 and TOM22 in the eluate as well as radiolabeled PINK1. Interestingly, we could also observe a faint band of PINK1 in the eluate when polarized organelles were used (Figure 1a). This finding demonstrates that the interactions of PINK1 with the TOM complex of healthy mitochondria is too weak to withstand the conditions of the BN-PAGE analysis.

**Figure 1.**
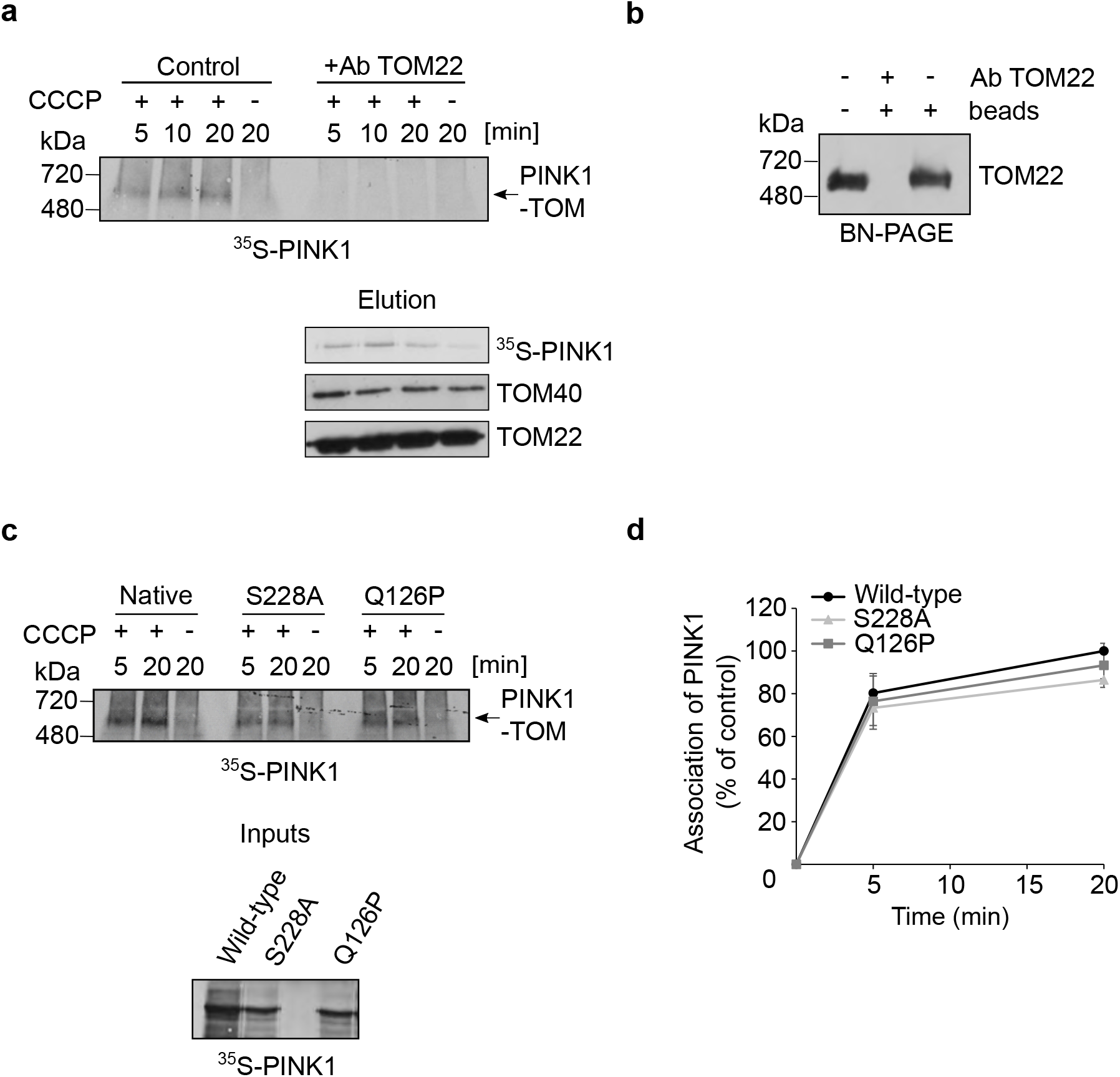
PINK1 associates with the TOM complex upon depolarization of mitochondria. **(a)** Radiolabeled PINK1 was imported into mitochondria for indicated time periods. In some cases, the isolated organelles were pretreated with CCCP. At the end of the import reactions, organelles were solubilized with digitonin-containing buffer and the indicated samples were subjected to pull-down with anti-TOM22 antibodies. Samples were then centrifuged and the supernatant was analyzed by BN-PAGE followed by autoradiography (upper panel). The PINK1-TOM complex species is indicated with an arrow. The immunodepleted proteins were analyzed by SDS-PAGE followed by autoradiography and western blotting using antibodies against TOM40 and TOM22 (lower panel). **(b)** Solubilized mitochondria were left untreated or were incubated with protein A beads in the presence or absence of TOM22 antibody. Samples were then analyzed by BN-PAGE followed by western blotting using TOM22 antibody. **(c)** Radiolabeled native PINK1 and the indicated mutants were incubated with isolated mitochondria. Further analysis was as described in panel (a). Lower panel: Radiolabeled samples that were used for the *in organello* import reactions were analyzed by SDS-PAGE and radiography. **(d)** The bands corresponding to PINK1 association with TOM complex in three independent experiments were quantified. The amount of assembled native PINK1 after 20 min was set to 100%.

To demonstrate the specificity of the TOM depletion, we confirmed that PINK1 could not be depleted from the 700 kDa species by antibodies against non-TOM components like Cytochrome C or VDAC1 (Suppl. Figure 1). We conclude that the formation of the 700 kDa species is a robust readout for stable association of PINK1 with the TOM complex. Next, we checked whether mutating an autophosphorylation site of PINK1 (S228A) or N-terminally localized glutamine residue, Q126P, a mutation reported in Parkinsonism, affect its association with the TOM complex. However, we did not observe any differences as compared to native PINK1 (Figure1 c, d). Our findings regarding the S228A variant are in line with previous findings and support the validity of our assay (Okatsu et al., 2013; Rasool et al., 2021). Thus, it appears that the defects related to these mutations do not arise from compromised interaction with the TOM complex.

### Identification of TOM70 as a potential receptor for PINK1 import into mitochondria

PINK1 is synthesized on cytosolic ribosomes and therefore, has to be targeted to mitochondria. Our first aim was to identify which receptors of the TOM complex could recognize PINK1 upon its translocation to the MOM. First, since we previously found TOM70 to play a role in PINK1 recognition (Kato *et al.*, 2013), we performed a peptide scan, in which peptides encompassing the N-terminal 140 amino acids of PINK1, a region that is sufficient for PINK1 targeting to mitochondria (Becker et al., 2012; Okatsu et al., 2015), were synthesized onto a membrane. Next, the membrane was incubated with purified recombinant cytosolic domain of yeast Tom70. In line with our previous observations, we detected interactions of the cytosolic domain of Tom70 with PINK1 peptides (Supplementary Figure 2a). We could observe Tom70 binding throughout PINK1 N-terminal sequence, with the highest intensity at MTS, OMS and TMD (Suppl. Figure 2b). These findings agree with current knowledge about the capacity of TOM70 to recognize hydrophobic internal segments of mitochondrial proteins as well as internal MTSs (Endo *et al.*, 2011; Backes *et al.*, 2018). To test the contribution of TOM70 in more detail, we used *in vitro* pull-down assay to study whether the cytosolic domains of Tom70 or Tom20 (for comparison) can bind radiolabeled PINK1. Our results revealed that whereas Tom20 has only a weak binding capacity to PINK1, Tom70 can do so with a much higher efficiency (Suppl. Figure 2c). Next, we were interested to find out whether the MTS in residues 1-35 of PINK1 is crucial to the recognition by TOM70. Of note, we could observe that both fragments of PINK1: 1-120 as well as 35-120 were bound by the Tom70 receptor suggesting that PINK1 recognition by Tom70 does not solely depend on its MTS (Suppl. Figure 2d). These results agree with previous reports (Zhou *et al.*, 2008; Okatsu *et al.*, 2015), which showed that PINK1 without its MTS could still be imported into depolarized mitochondria. Thus, we concluded that TOM70 could be the primary receptor for PINK1 import into mitochondria by recognizing the OMS and the TMD elements of the protein.

**Figure 2.**
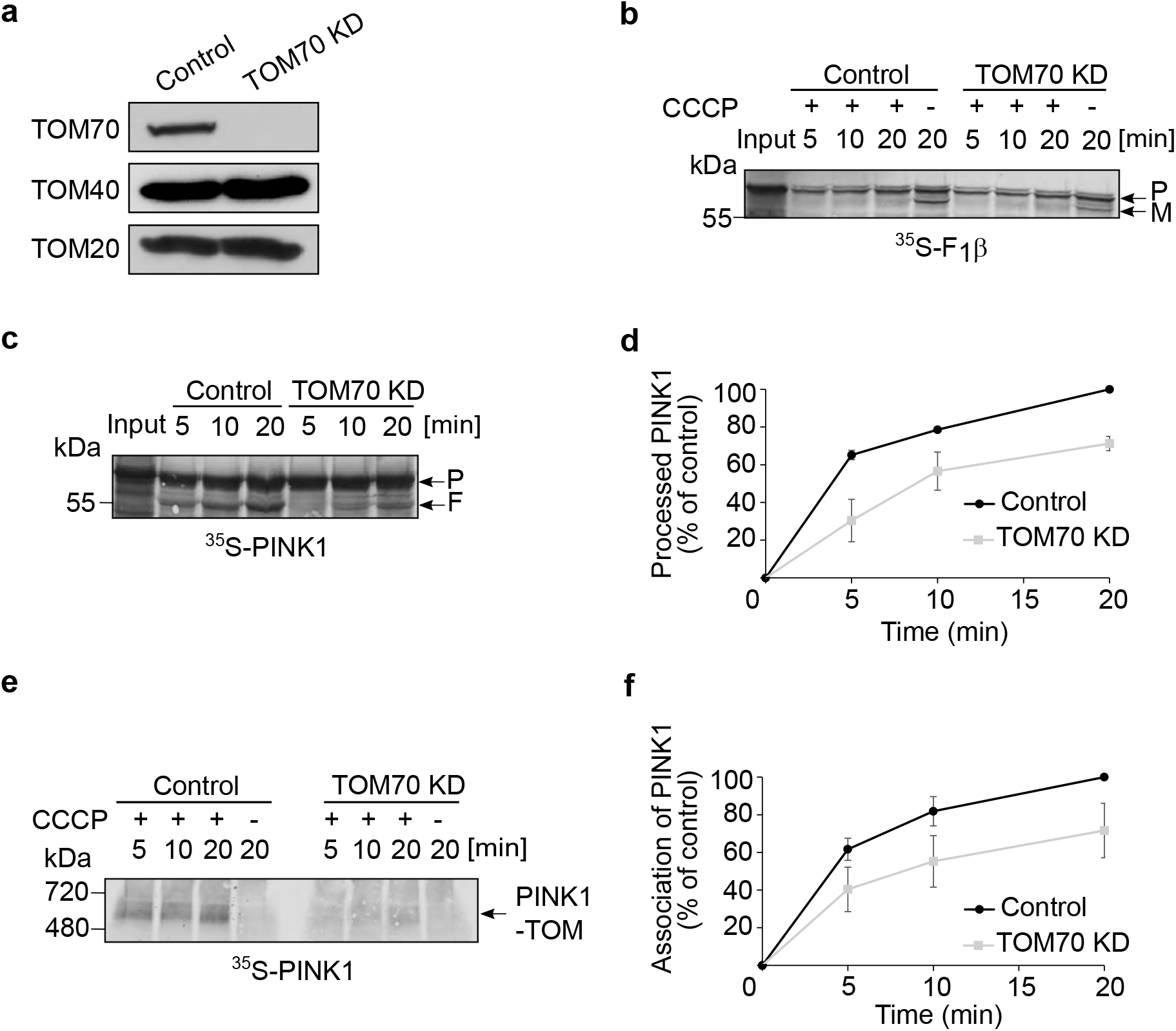
TOM70 plays an important role in PINK1 import into mitochondria. **(a)** Mitochondria were isolated from either control or cells knocked down for TOM70. Isolated organelles (50 μg) were analyzed by SDS-PAGE and western blotting with the indicated antibodies. **(b)** Radiolabeled F_1_β was incubated for the indicated time periods with mitochondria isolated from either control or TOM70 KD cells in the absence or presence of CCCP. Subsequently, samples were analyzed by SDS-PAGE and autoradiography. The precursor (P) and mature (M) forms of the protein are indicated. **(c)**. Radiolabeled PINK1 was incubated with mitochondria isolated from either control or TOM70 KD cells. Next, samples were analyzed by SDS-PAGE and autoradiography. The bands corresponding to the precursor form of PINK1 (P) or the PARL-processed form (F) are indicated. **(d)** The bands corresponding to processed PINK1 (F) were quantified and the intensity of the band corresponding to import for 20 min into control organelles was set to 100%. The results present the average of three independent experiments (± SD). **(e)** Import into the indicated isolated organelles was performed and analyzed as described in the legend to Fig. 1a. **(f)** The bands corresponding to PINK1 association with TOM complex were quantified. The intensity of the band corresponding to assembled PINK1 in control mitochondria after 20 min was set up to 100%. The results represent the average of three independent experiments (± SD).

### TOM70 is important for PINK1 import into mitochondria

To further validate the importance of TOM70 receptor, we knocked it down in HeLa cells and isolated mitochondria from either control or these manipulated cells. Figure 2a shows the validation by western blotting of the TOM70 depletion in the isolated organelles. To exclude general problem in the import capacity of mitochondria isolated from the TOM70 KD cells, we imported into these organelles radiolabelled F1β (subunit of ATP synthase), which requires the presence of ΔΨm for its translocation to the matrix where it is processed from the precursor (P) form to the mature (M) one by removal of the MTS (Figure 2b, lanes with no CCCP addition). As expected for an MTS-containing matrix substrate (Backes et al., 2018), the import of this precursor protein into mitochondria depleted for TOM70 was comparable to its import into control organelles (Figure 2b). In contrast, when radiolabeled PINK1 was imported into control and TOM70 KD mitochondria that were either polarized (Figure 2c) or depolarized by addition of CCCP (Figure 2e), the absence of TOM70 resulted in a significant retardation of the import capacity of the organelles. Import into healthy organelles was measured by the appearance of 52 kDa-processed version of PINK1 (F, Figure 2c, d) and association of PINK1 with the TOM complex in depolarized mitochondria was quantified based on the built-up of the 700 kDa species upon BN-PAGE analysis (Figure 2e, f). In both cases, PINK1 import was reduced to about 70% of the control. Hence, we propose that TOM70 plays a central role in PINK1 recognition at the MOM and subsequent import into both healthy and depolarized mitochondria.

### TOM20 plays only a minor role in PINK1 recognition at the MOM

Our in vitro binding experiments described in Suppl. Figure 2c suggested that TOM20 has only a weak binding capacity to PINK1. To check whether TOM20 could play a role in PINK1 recognition *in vivo*, we depleted U2OS cells of the TOM20 receptor and isolated mitochondria from these cells (Figure 3a). Initially, we controlled whether MTS containing precursor protein like F1β is affected by the absence of TOM20. As expected, we observed a reduction in the import capacity of this protein into organelles lacking TOM20 and in both types of organelles the import was eliminated by adding CCCP (Figure 3b). Next, we imported radiolabeled PINK1 into polarized mitochondria isolated from either control or TOM20 KD cells and found that PINK1 processing by PARL was hardly affected, if at all, by the absence of TOM20 (Figure 3c, d). Along the same line, association of PINK1 with the TOM complex in depolarized mitochondria was decreased in the TOM20 depleted organelles by only 20% (Figure 3e, f). Thus, we concluded that TOM20 does play, however rather marginal, a role in PINK1 recognition.

**Figure 3.**
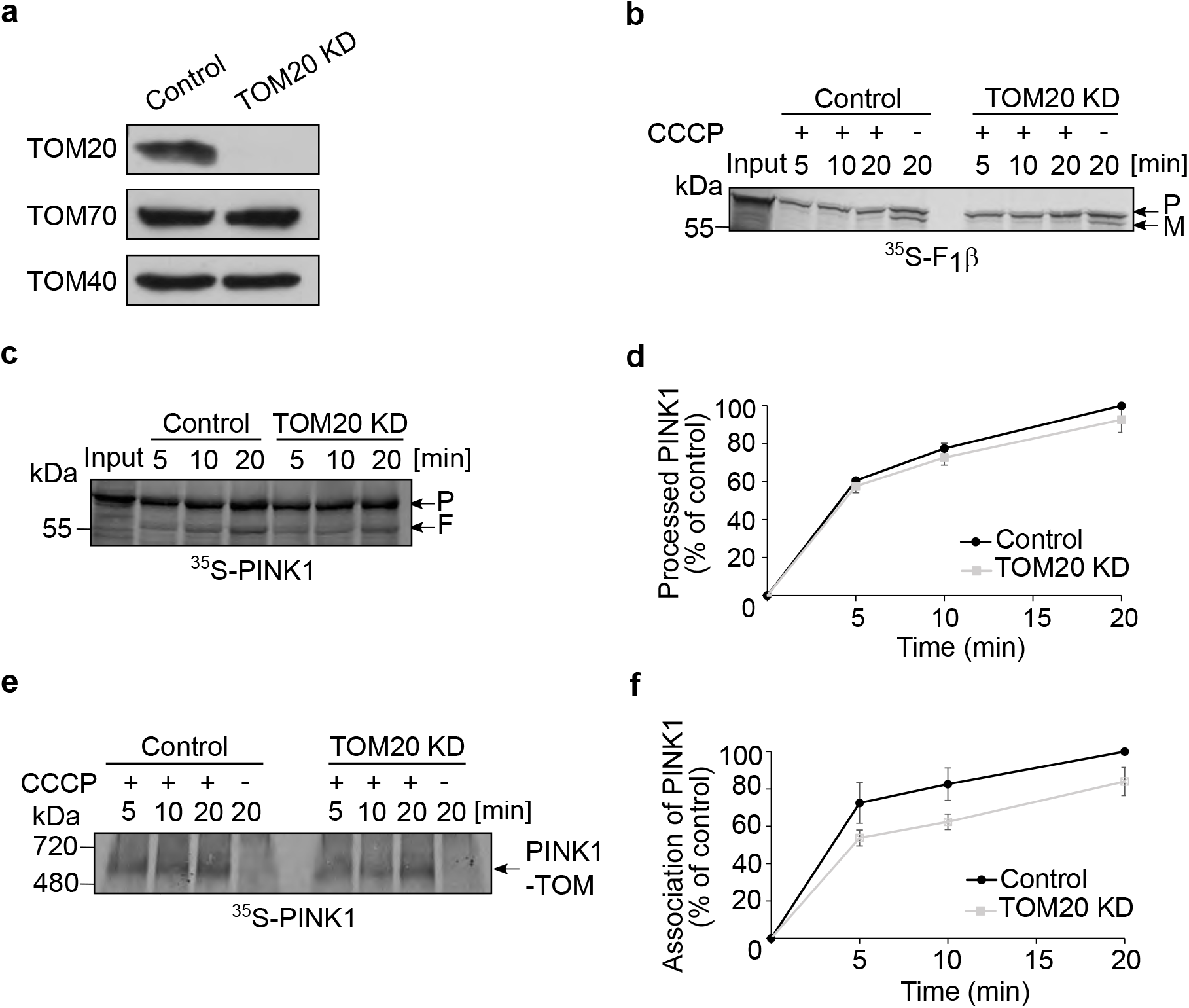
TOM20 plays only a minor part in PINK1 recognition at the MOM. **(a)** Mitochondria were isolated from either control or cells knocked down for TOM20. Isolated organelles (50 μg) were analyzed by SDS-PAGE and western blotting with the indicated antibodies. **(b)** Radiolabeled F_1_β was imported and analyzed as described in the legend to Fig. 2b. **(c, d)** Radiolabeled PINK1 was incubated with mitochondria isolated from either control or TOM20 KD cells. Further analysis and quantification were as described in the legends to Fig. 2c, d. **(e, f)** Import into the indicated isolated organelles was performed and analyzed as described in the legend to Fig. 1a. Quantification of bands was as described in the legend to Fig. 2f.

### Double knockdown of TOM70 and TOM20 has a synthetic effect on PINK1 import

Since we observed that PINK1 is imported into mitochondria even upon the individual knock down of the import receptors, we wondered whether in the absence of one receptor, the other one can take-over its function. To address this possibility, we knocked down both proteins in HeLa cells and then isolated mitochondria. While TOM70 was not detected at all in the organelles from the double KD cells, we detected faint band of TOM20 in these cells (Figure 4a). Of note, the levels of the central TOM subunit, TOM40 was not affected by the practical absence of both receptors. As previously, we used F_1_β import as a control substrate and as an indication for successful CCCP-treatment (Figure 4b). Indeed, while certain levels of the protein were imported into organelles lacking either TOM70 (Fig. 2b) or TOM20 (Fig. 3b), the removal of both receptors eliminated completely the import of F_1_β (Figure 4b). In sharp contrast, when radiolabeled PINK1 was imported into control and double KD polarized mitochondria the processing of PINK1 by PARL remained at approximately 80% (Figure 4c, d). The dependency on the receptors was different when depolarized organelles were used. Under these conditions, association of PINK1 with the TOM complex lacking both receptors dropped to approximately 40% (Figure 4e, f)., These findings support our assumption that basically, both receptors can recognize PINK1 and in the absence of a single receptor, the other can perform most of the recognition. Next, we investigated the possibility that secondary effects of the double depletions on the fitness of the TOM complex caused the dramatic reduction in the import of PINK1. Monitoring the assembly of the TOM complex by BN-PAGE revealed that the migration of the translocase from the double depleted cells remained unchanged (Figure 4e, lower panel). This observation agrees with previous findings that the peripheral receptors TOM20 and TOM70 are not part of the TOM core complex, which is detected by BN-PAGE under these conditions. Collectively, we suggest that the strong effect of the double depletion of TOM70 and TOM20 indicates the potential exchangeable roles of both receptors in protein recognition. The fact that double knockdown only moderately affected PINK1 import and PARL-processing in polarized mitochondria and on the other hand, had a drastic effect on association with the TOM complex of depolarized mitochondria, might suggest that both receptors have a stabilizing role in the association of PINK1 with the TOM complex.

**Figure 4.**
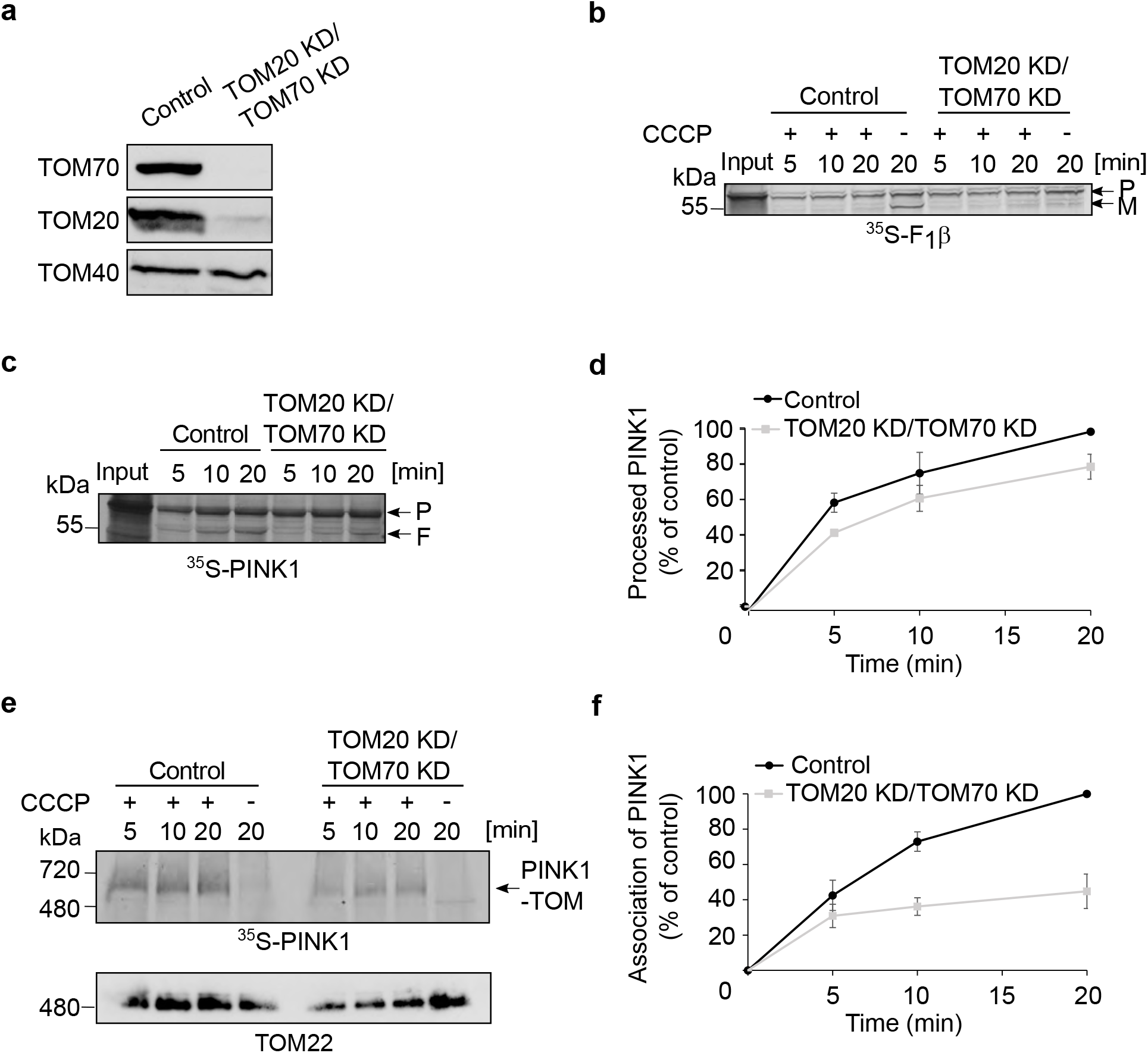
Double knockdown of both receptors TOM70 and TOM20 compromises the import of PINK1 into depolarized mitochondria. **(a)** Mitochondria were isolated from either control or cells knocked down for both TOM70 and TOM20. Isolated organelles (50 μg) were analyzed by SDS-PAGE and western blotting with the indicated antibodies. **(b)** Radiolabeled F_1_β was imported and analyzed as described in the legend to Fig. 2b. **(c, d)** Radiolabeled PINK1 was incubated with mitochondria isolated from either control or TOM20 KD/TOM70 KD cells. Further analysis and quantification were as described in the legends to Fig. 2c, d. **(e, f)** Import into the indicated isolated organelles was performed and analyzed as described in the legend to Fig. 1a. Quantification of bands was as described in the legend to Fig. 2f. The lower panel of (e) is an immunodecoration of the same membrane with antibodies against TOM22.

### PINK1 association with the TOM complex is reduced upon KD of TOM7 and TOM7/TOM70

TOM7 was found to play a role in the integration of PINK1 into the MOM (Hasson *et al.*, 2013; Sekine *et al.*, 2019). However, it is still unclear how TOM7 facilitates this process. To test for potential involvement of TOM7 in the association with the TOM complex, we isolated mitochondria from TOM7 KD U2OS cells. Since we could not find a functional antibody against TOM7, we verified the KD by performing a RT-qPCR and detecting dramatically lower levels of the encoding mRNA (not shown). To substantiate the assumption that the levels of TOM7 are indeed profoundly reduced, we followed the behaviour of the TOM complex on BN-PAGE. It was previously reported that upon TOM7 depletion, the levels of the TOM complex are decreased, and the complex migrates at apparently lower molecular mass (Kato & Mihara, 2008). In agreement with this previous report, the TOM complex in the samples from the TOM7 KD cells was detected as a weaker band, which migrated faster than the control complex (Figure 5a). Moreover, when radiolabeled TOM40 was imported *in organello* into mitochondria isolated from the KD cells, it integrated into a smaller species of the TOM complex as compared to the species observed with organelles isolated from untreated cells or those that obtained scrambled siRNA (Figure 5b). Immunodecoration of the same membrane with antibodies against TOM40 confirmed this observation (Figure 5b, lower panel). Taken together, we concluded that TOM7 was indeed depleted in these KD cells.

**Figure 5.**
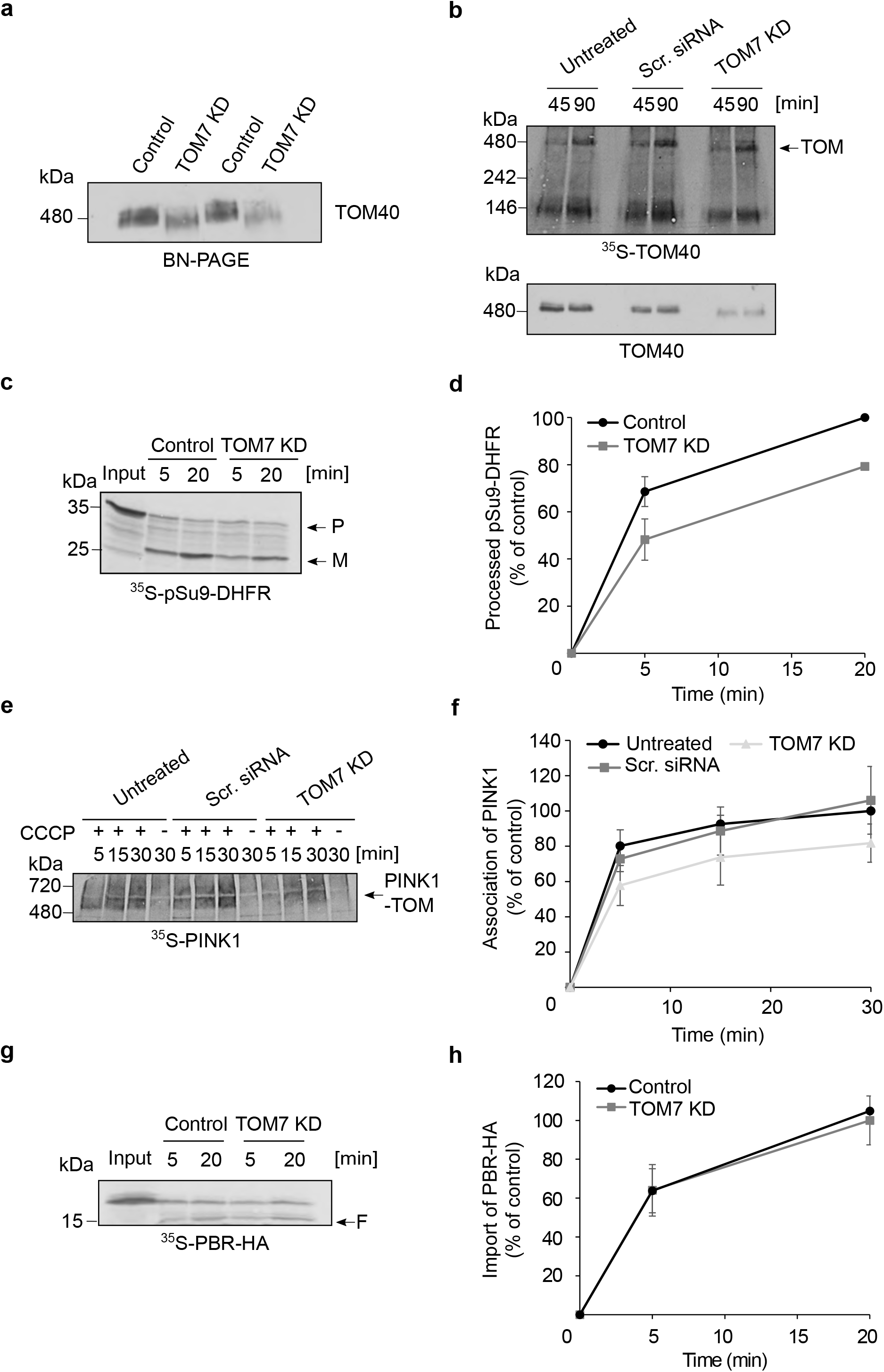
PINK1 association with the TOM complex is reduced upon depletion of TOM7. (a) **Mitochondria isolated from control or TOM7 KD cells were analyzed by** BN-PAGE followed by immunodecoration with an antibody against TOM40. Duplicates of each sample were loaded in an alternating manner to emphasize the different levels and migration behavior of the TOM complex. **(b)** Mitochondria were isolated from untreated U2OS cells or from cells treated with either scrambled (Scr.) siRNA or TOM7 siRNA. Radiolabeled TOM40 was imported into the isolated organelles for either 45 or 90 minutes and samples were analyzed via BN-PAGE and autoradiography (upper panel). After exposing to autoradiography, the same membrane was immunodecorated with TOM40 antibody to visualize the TOM endogenous complex (lower panel). The assembled TOM complex is indicated with an arrow. **(c)** Radiolabeled pSu9-DHFR was imported into control and TOM7 KD mitochondria for the indicated time periods. Unimported proteins were removed with PK and samples were analyzed via SDS-PAGE and autoradiography. The precursor (P) and mature (M) forms are indicated. **(d)** The bands corresponding to processed pSu9-DHFR were quantified and the intensity of the band corresponding to import into control mitochondria for 20 min was set to 100%. The results represent the average of three independent experiments (± SD). **(e, f)** Import into the indicated isolated organelles was performed and analyzed as described in the legend to Fig. 1a. Quantification of bands was as described in the legend to Fig. 2f. **(g)** Radiolabeled PBR was incubated with either control or TOM7 KD mitochondria for the indicated time periods. PK was added at the end of the import reaction and samples were then analyzed by SDS-PAGE and detected via autoradiography. A typical proteolytic fragment of membrane integrated PBR is indicated (F). **(h)** Bands representing the proteolytic fragment of PBR were quantified. the intensity of the band corresponding to import into control mitochondria for 20 min was set to 100%. The results represent the average of three independent experiments (± SD).

Subsequently, we wanted to check how the absence of TOM7 would affect import of matrix residing protein. The import of pSu9-DHFR, as monitored by its processing by MPP, was compromised by 20-30% upon the depletion of TOM7 (Figure 5c, d), probably due to the reduced stability of the TOM complex. A similar degree of reduction was also observed upon studying the association of PINK1 with the TOM complex of depolarized organelles from the TOM7 KD cells (Figure 5e, f). Thus, it seems that TOM7 depletion does not specifically affect PINK1 MOM integration, but also other proteins that depend on the correct conformation of the TOM complex for their import. To corroborate this suggestion, we imported a multi-span MOM protein, PBR (TPSO), that was previously reported to insert in the MOM independently of the TOM complex (Otera *et al.*, 2007). Indeed, TOM7 KD did not affect PBR integration into the MOM (Figure 5g, h).

Next, we aimed to investigate whether double depletion of TOM7 and TOM70 would have a synthetic effect on PINK1 association with mitochondria. TOM7O KD was confirmed by immunodecoration (Figure 6a), whereas the depletion of TOM7 via the altered migration behaviour of the TOM complex in BN-PAGE (Figure 6b, lower panel). When radiolabeled PINK1 was imported into TOM7/TOM70 double depleted depolarized mitochondria, we observed a dramatic reduction of more than 60% in the association of PINK1 to the TOM complex as compared to control organelle (Figure 6b). This striking reduction is probably the outcome of the combined effects of absence of the main receptor for PINK1 import, TOM70, as well as the destabilization of the TOM complex in the absence of TOM7.

**Figure 6.**
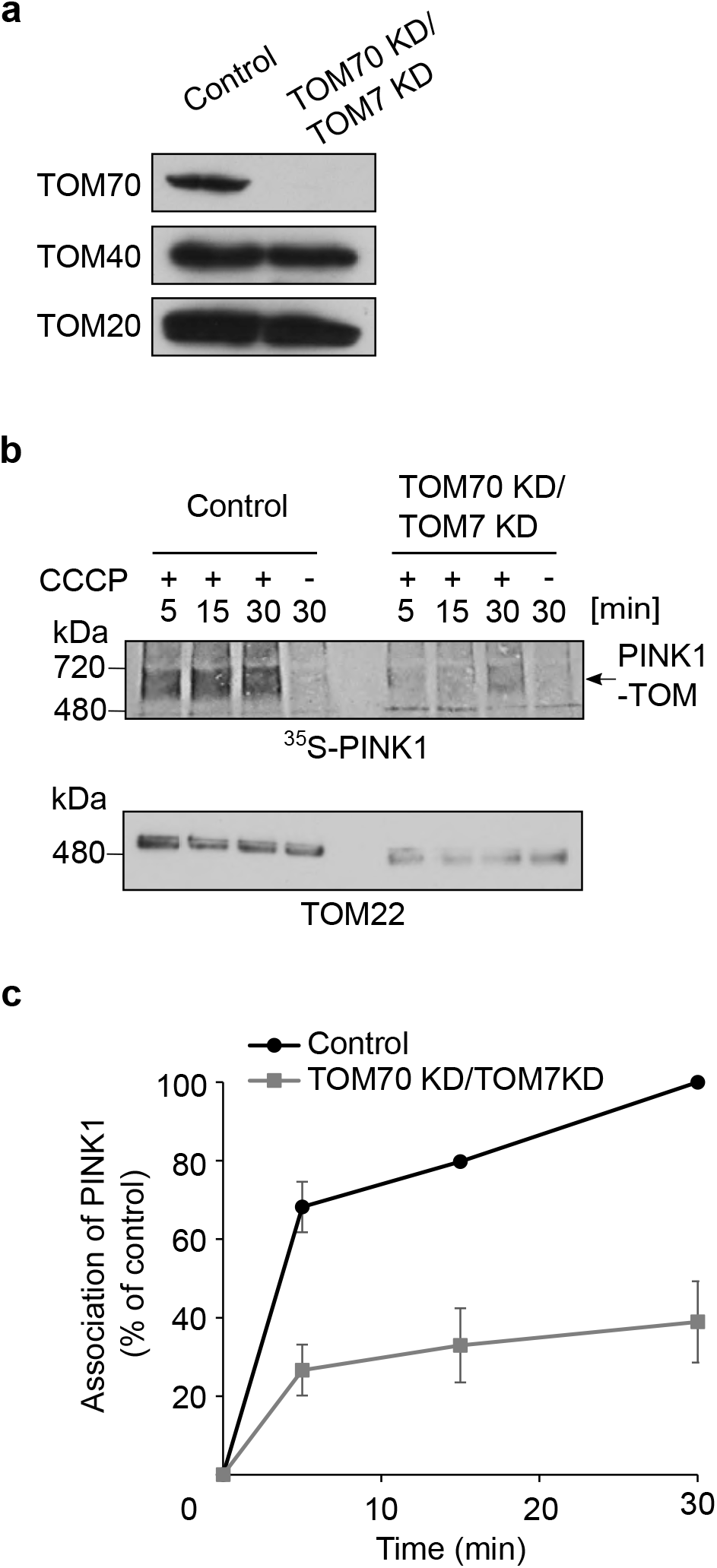
Double knockdown of TOM70 and TOM7 has a dramatic effect on PINK1 import into depolarized mitochondria. **(a)** Mitochondria were isolated from either control or cells knocked down for both TOM70 and TOM7. Isolated organelles (50 μg) were analyzed by SDS-PAGE and western blotting with the indicated antibodies. **(b)** Radiolabeled PINK1 was imported into either control or TOM7/TOM70 KD depolarized mitochondria for the indicated time periods. Samples were subsequently analyzed by BN-PAGE and autoradiography (upper panel). After autoradiography exposures, the same membrane was incubated with TOM22 antibody to show the appearance of the TOM complex (lower panel). **(c)** The bands corresponding to PINK1 association with the TOM complex were quantified. The band corresponding to assembled PINK1 after 20 min import into control mitochondria was set to 100%. The results represent the average of three independent experiments (± SD).

### The lumen of the TOM pore is crucial for PINK1 association with the MOM

There are three main hypotheses in the literature regarding the precise location of PINK1 in the MOM of depolarized organelles.: The first suggests that PINK1 is laterally released from the TOM complex into the bulk of the outer membrane (Nguyen et al., 2016), the second argues that PINK1 is arrested in the TOM40 lumen from where it can trigger the mitophagy cascade (Hasson *et al.*, 2013), whereas the third proposes that the N-terminal segment of PINK1 could undergo lateral release, but remains bound to the TOM complex and is not released into the bulk OM (Rasool et al., 2021). Since all alternatives emphasize the role of TOM40 in this process, we aimed to study the dependence of PINK1 on the TOM pore upon import into CCCP-treated mitochondria. TOM40 is the central essential component of the TOM complex and therefore its knockdown could lead to many secondary effects such as altered levels of many additional other mitochondrial proteins. Thus, we used an alternative method namely, the addition of excess amounts of purified recombinant pSu9-DHFR to clog the TOM complex. Ideally, the MTS of Su9 would drive the protein into the matrix, however by addition of the DHFR ligand, methotrexate, DHFR domain becomes tightly folded and disables the translocation of the protein across the TOM complex. The latter results in the blockage of the TOM entry gate, making it almost impossible for other proteins to pass through it. As expected, import of PINK1 and F_1_β into polarized mitochondria that were blocked by pSu9-DHFR was acutely affected (Figure 7a and b, respectively). Both proteins are translocated into the matrix via the presequence pathway, therefore they rely on TOM40 as their initial entry gate.

**Figure 7.**
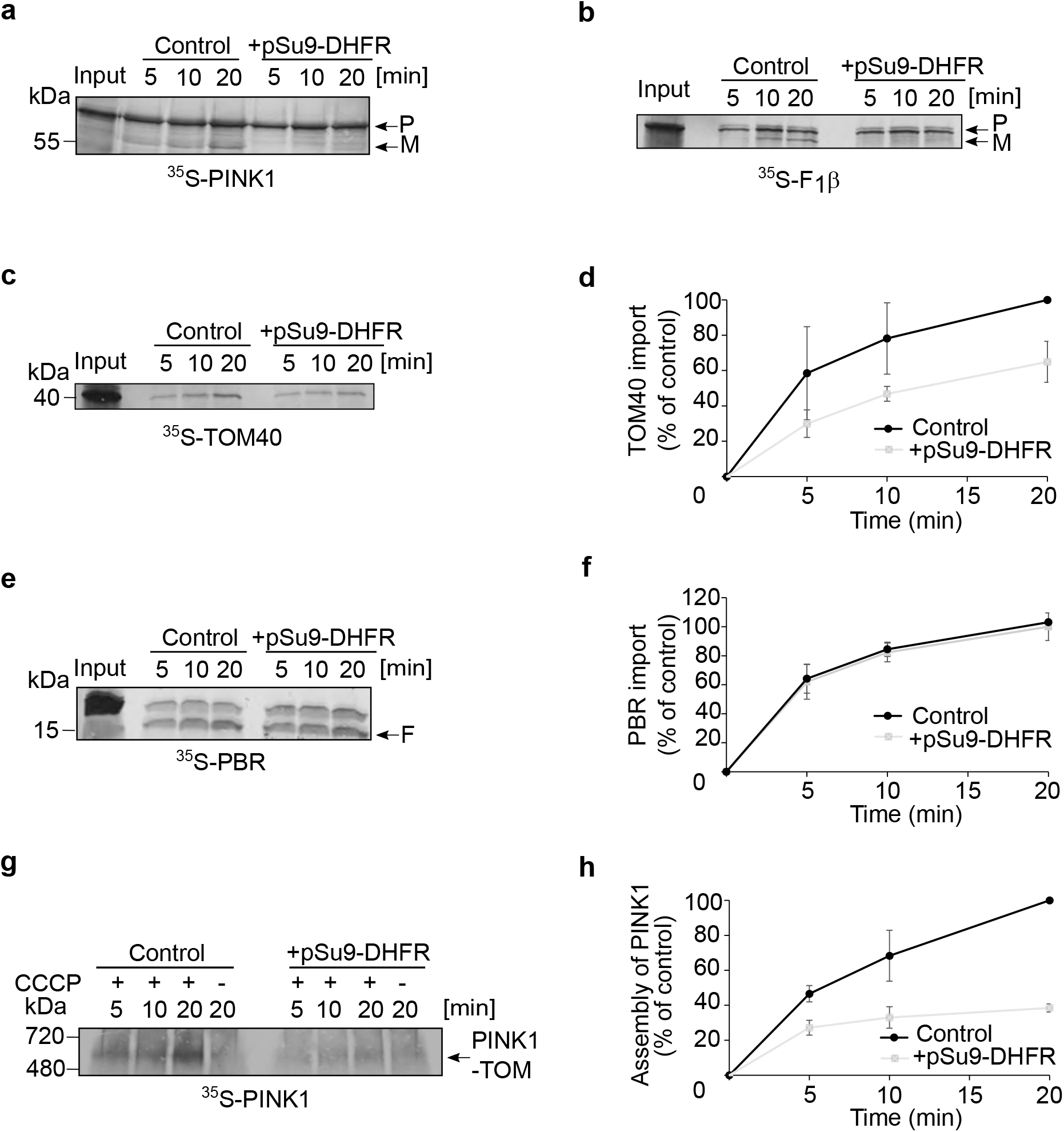
TOM40 lumen is crucial for PINK1 association with the TOM complex. Isolated polarized mitochondria were preincubated in the absence or presence of purified pSu9-DHFR for 10 min on ice prior to the import of the radiolabeled proteins. Radiolabeled PINK1 **(a)**, F1β **(b)**, or TOM40 **(c)** were imported into the isolated organelles Samples were then analyzed by SDS-PAGE followed by autoradiography. **(d)** The bands corresponding to membrane integrated TOM40 were quantified. The band corresponding to integrated TOM40 after 20 min import into untreated mitochondria was set to 100%. The results represent the average of three independent experiments (± SD). (e) Radiolabeled PBR was incubated with either treated or untreated mitochondria for the indicated time periods. PK was added at the end of the import reaction and samples were then analyzed by SDS-PAGE and detected via autoradiography. A typical proteolytic fragment of membrane integrated PBR is indicated (F). **(f)** Bands representing the proteolytic fragment of PBR were quantified. the intensity of the band corresponding to import into control mitochondria for 20 min was set to 100%. The results represent the average of three independent experiments (± SD). **(g)** Radiolabeled PINK1 was imported into either control or pSu9-DHFR treated mitochondria for the indicated time periods. Samples were subsequently analyzed by BN-PAGE and autoradiography. **(h)** The bands corresponding to PINK1 association with the TOM complex were quantified. The band corresponding to assembled PINK1 after 20 min import into control mitochondria was set to 100%. The results represent the average of three independent experiments (± SD).

Along the same line, and in agreement with previous results with fungal organelles (Rapaport and Neupert, 1999), when radiolabeled TOM40 was imported, its integration into the MOM was reduced to around 60% when the entry channel was clogged with pSu9-DHFR (Figure 7 c, d). This reduction was more moderate than the one observed for the matrix proteins because the biogenesis of TOM40 does not require TIM23 and depends on the TOM and SAM complexes that are more abundant than the TOM/TIM23 supra-complexes. As expected, when PBR, which is not dependent on the TOM complex for its membrane assembly, was imported into mitochondria preincubated with pSu9-DHFR, we could not observe any difference as compared to control conditions (Figure 7e, f). This observation suggests that there is no general defect in the import capacity of mitochondria upon the addition of the clogger. Finally, we imported radiolabelled PINK1 into depolarized mitochondria in which TOM40 was blocked by pSu9-DHFR. We observed a dramatic decrease in the PINK1 association with the TOM complex to less than 40% as compared to control reactions (Figure 7g, h). Thus, these experiments strongly suggest that PINK1 uses the lumen of TOM40 for its stable association with the TOM complex.

### TcPINK1 apparently blocks the TOM pore in depolarized mitochondria

Lastly, we aimed to determine which scenario is more probable upon mitochondrial depolarization: PINK1 is laterally released into the bulk of the MOM, gets stalled in the TOM40 pore, or remains bound to TOM, but outside the TOM40 barrel. To address this question, we purified recombinant TcPINK1, which was previously used in studies on structure-function relationships of PINK1 (Kumar *et al.*, 2017; Rasool *et al.*, 2018; Okatsu *et al.*, 2018; Rasool et al., 2021). We opted for TcPINK1 rather that human PINK1 since the former is more stable and can be easier purified as a recombinant protein. Initially, we confirmed that TcPINK1 can associate in depolarized mitochondria with human TOM complex to the same oligomeric species as human PINK1 (hPINK1) (Figure 8a). Next, we performed reciprocal experiments to the ones described above by testing whether TcPINK1 can clog the TOM complex and thereby inhibit the import of other TOM-dependent substrates. Hence, we performed *in organello* import assays in which mitochondria were first depolarized and incubated with TcPINK1. Subsequently, radiolabeled TOM40 or hPINK1 were imported into these treated organelles. We observed that both proteins were compromised in their integration into the MOM under these conditions (Figure 8b-e). Of note, import of TOM40 into mitochondria in which the TOM pore was blocked with either pSu9-DHFR (Figure 7c, d) or TcPINK1 (Figure 8b, c) resulted in a similar import reduction to approximately 60%. Although, we cannot exclude the possibility that the kinase domain of PINK1 contributes to the import inhibition by hindering access of other substrates to the TOM pore, we assume that most of the competition is because PINK1, similarly to pSu9-DHFR, occupies the lumen of the TOM pore in depolarized mitochondria.

**Figure 8.**
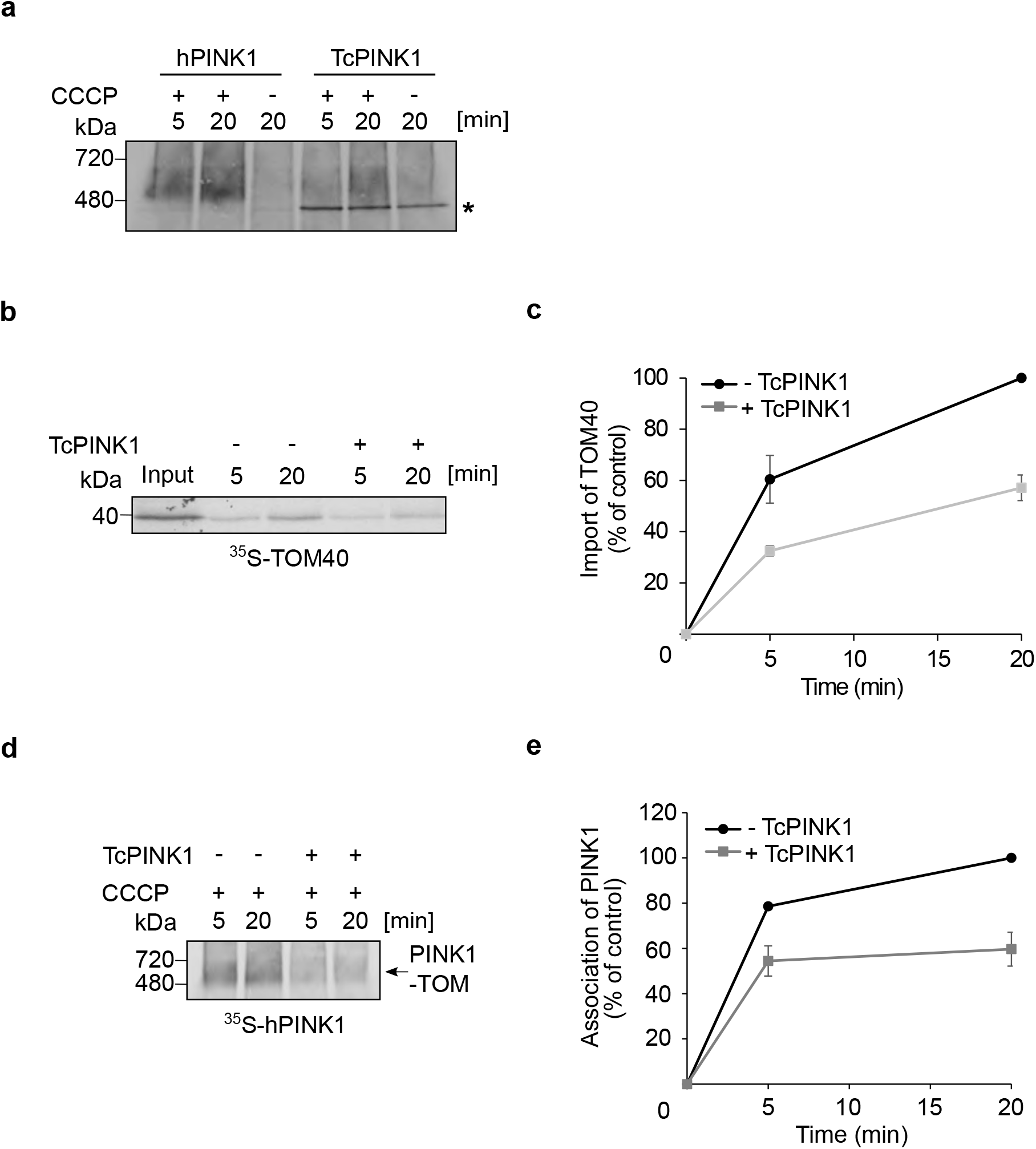
TcPINK1 apparently blocks the TOM pore in depolarized mitochondria. **(a)** Radiolabeled hPINK1 and TcPINK1 were imported into mitochondria isolated from U2OS cells. The organelles were depolarized prior to the import in the indicated samples. Samples were then analyzed by BN-PAGE and autoradiography. *, indicates an unspecific band. **(b)** Radiolabeled TOM40 was incubated for indicated time periods with depolarized mitochondria that were preincubated in the presence or absence of recombinant TcPINK1. At the end of the import reactions sample were subjected to alkaline extraction and the pellet fractions were analyzed by SDS-PAGE and autoradiography. **(c)** The bands corresponding to membrane-embedded TOM40 were quantified. The band corresponding to membrane embedded TOM40 after 20 min import into control mitochondria was set to 100%. The results present the average of two independent experiments (± SD). **(d)** Radiolabeled hPINK1 was incubated for indicated time periods with depolarized mitochondria that were preincubated in the presence or absence of recombinant TcPINK1. At the end of the import reactions, sample were analyzed by SDS-PAGE and autoradiography. (**e**) The bands corresponding to PINK1 association with the TOM complex were quantified. The intensity of the band corresponding to assembled PINK1 in control mitochondria after 20 min was set up

## Discussion

PINK1 import into healthy mitochondria is relatively well described, however its fate in depolarized organelle remained unclear. In the current study, TOM70 was found to be the main receptor for PINK1 import into either healthy or compromised mitochondria. Our results are in line with previous report (Kato *et al.*, 2013) and demonstrate in addition direct interaction between both proteins. Although the single depletion of TOM20 did not result in a major reduction of PINK1 interactions with the TOM complex, the double KD of both receptors TOM20 and TOM70, caused a dramatic decrease in this interaction, which was more than expected according to the additive effects of the single KDs. Thus, we conclude that, although TOM70 plays the primary role, both receptors can contribute to the import of PINK1 and if one of them is missing, the other can compensate (at least partially) for this absence. The fact that PINK1 contains a classical MTS and yet does not primarily depend on TOM20, shows that either, as reported before for other substrates (Backes et al. 2018), TOM70 can also recognize this MTS, or that the internal signals like the OMS or the TMD, which can be recognized by TOM70, are playing a superior role in recognition at the surface of the organelle under these conditions. Of note, our peptides scan assay showed that TOM70 can bind all the three elements (MTS, OMS, and TMD) of PINK1 and thus the two aforementioned options are not mutually exclusive.

Regardless of the identity of the primary receptor, after the initial recognition, PINK1 molecules are relayed to further locations within the MOM of depolarized organelles. A previous report on that topic suggested that TOM7 mediates the lateral translocation of the PINK1 from the TOM40 pore (Sekine *et al.*, 2019). However, the recently published atomic structure of the human TOM complex, where the β-barrels formed by TOM40 appear to be tightly closed (Wang et al., 2020; Guan et al., 2021), raised the question if such a lateral release can be anticipated. As was already published before (Kato and Mihara, 2008), our findings suggest that the TOM complex in the absence of TOM7 is less stable. If TOM7 is indeed a gate for a lateral release, we would assume that initial PINK1 association with the TOM complex should not be affected, but rather further downstream steps. However, we observe that the absence of TOM7 causes a reduction in the initial association of PINK1with the TOM complex. Therefore, and considering the observation that also other substrates are affected by depletion of TOM7, we propose that the contribution of TOM7 to the association of PINK1 with the MOM is probably rather indirect and occurs through the stabilization effect of this protein on the overall structure of the TOM complex. On the other hand, we cannot exclude that PINK1 binds both TOM7 and TOM70 at two different places, which would explain why the double KD TOM7/TOM70 has such a dramatic effect on formation of PINK1-TOM.

In an additional set of experiments, we demonstrate that PINK1 association with the TOM complex is compromised by clogging the lumen of this complex and reciprocally, PINK1 itself can block the import pore of the translocase. Hence, although previous reports suggested a PINK1 lateral release from the TOM complex (Lazarou *et al.*, 2012; Sekine *et al.*, 2019), we favour the possibility that PINK1 remains at the lumen of the TOM40 in depolarized organelles. The following arguments support this proposal: (i) PINK1 co-migrates with the core TOM complex on BN-PAGE that consists of TOM40, TOM22, and accessory small TOMs: TOM5, TOM6, TOM7. This means that the interactions of PINK1 with the TOM complex are stable enough to sustain the analysis conditions. (ii) Blocking of the TOM channel by PINK1 would also prevent an import of all other TOM-dependent substrate proteins. Such a blockage would be prudent as the malfunctional compromised organelles are destined for degradation upon induction of mitophagy. (iii) PINK1 was reported to be reimported and cleaved by PARL rather fast upon re-polarization of mitochondria (Lazarou *et al.*, 2012). Location of PINK1 at the pore of the TOM complex can readily explain such a fast re-import. In contrast, re-translocation from the lipid core of the MOM would require initially an extraction from the membrane and then, a putative re-entering route into the TOM complex. Nevertheless, despite these supportive arguments, only future structural studies of the PINK1-TOM adduct would be able to resolve this issue. In summary, our current study provides new insights into the crucial initial stage of PINK1-dependent mitophagy namely, the anchoring of PINK1 to the MOM of compromised mitochondria.

## Materials and Methods

### Cell culture and induction of knockdown of selected proteins

HeLa and U2OS cells were used in the current study, and both were cultured at 37°C under 5% CO_2_ in Dulbecco’s modified Eagles medium (DMEM, Gibco) supplemented with 10% (vol/vol) fetal bovine serum (FBS). To knock down TOM70, HeLa cells expressing shRNA against TOM70 under the control of doxycycline promoter were used (Kozjak-Pavlovic et al., 2007). To deplete TOM70, 1 μg/ml doxycycline was added every second day for 7 days.

TOM20 and TOM7 KDs were induced with FlexiTube siRNA purchased from QIAGEN (SI00301959 and SI04364955, respectively). The target sequences were 5′-AAAGTTACCTGACCTTAAAGA-3’ in the case of TOM20 and 5’-CACGGCCGTCGCCATGGTGAA-3’ for TOM7. U2OS cells were transfected twice with these siRNAs with a 24 h interval. After 72 h, the cells were harvested, and mitochondria isolated. To create TOM70/TOM20 or TOM70/TOM7 double KD lines, HeLa cells were first induced with 1 μg/ml doxycycline for 24 h and then transfected with siRNA against either TOM20 or TOM7 twice within 24 h interval. Each time the cells were recovered with DMEM containing 20% FBS and 2 μg/ml doxycycline. Cells were harvested 72 hours after the second transfection with TOM20 or TOM7 siRNA, and isolation of mitochondria was performed. According to the required amount of isolated mitochondria, cells were seeded in different numbers of TC Dish 150 Standard plates and grown until 80% confluent.

### Isolation of mitochondria

Cells were washed once with PBS buffer and scraped from the plate using a spatula before transferring the cell suspension to a small test tube. Subsequently, the cells were centrifuged (300 x g, 5 min, 4°C) and resuspended in HMS-A buffer (0.22 M mannitol, 0.07 M sucrose, 0.02 M HEPES-KOH, pH 7.6, 1 mM EDTA, 0.1% BSA, 1 mM phenylmethylsulfonyl fluoride (PMSF)). The used buffers were described previously (Becker *et al.*, 2012). Afterwards, the samples were passed nine times through needles of different sizes (20G, 23G, and 27G, Sterican). The homogenized lysates were centrifuged (900 x g, 5 min, 4°C) and the supernatants were collected whereas the pellets were discarded. The latter supernatants were centrifuged (9000 x g, 15 min, 4°C) to obtain crude mitochondria in pellets. The pellets were then washed with HMS-B buffer (HMS-A buffer lacking BSA) and centrifuged again (10000 x g, 10 min, 4°C). The pellets representing the mitochondrial fraction were used for *in vitro* import assays.

### Mitochondrial*in vitro* import followed by SDS-PAGE or BN-PAGE

Radiolabeled proteins were synthesized in the presence of ^35^S-methionine and ^35^S-cysteine using either coupled reticulocyte lysate system (TNT, Promega) or uncoupled reticulocyte lysate system (Promega). For the import reaction, 50-100 μg of isolated mitochondria were resuspended in HS buffer (20 mM HEPES-KOH, pH 7.6, 250 mM sucrose, 5 mM magnesium acetate, 80 mM potassium acetate, 7.5 mM glutamate, 5 mM malate, 1 mM DTT) and incubated at 30°C for different time periods with radiolabeled proteins in the presence of 4 mM ATP and 2 mM NADH. Where indicated, mitochondria were pre-incubated for 5 min on ice with 20 μM of the uncoupler carbonyl cyanide chlorophenylhydrazone (CCCP) prior to the import reactions.

To clog the TOM pore, isolated organelles were preincubated on ice for 10 min with 20 μg of recombinantly expressed pSu9-DHFR-His, in the presence of 1 mM methotrexate and 1 mM NADPH prior to the import of the radiolabeled proteins. Alternatively, the TOM pore was blocked by addition of 10-20 μg of TcPINK1 for 10 min at 30°C. At the end of the import reactions, samples were centrifuged (13200 x g, 10 min, 4°C) and handled differently depending on the expected localization of the protein of interest within the organelle. Isolated mitochondria with imported outer membrane proteins such as PBR/TSPO and matrix localized proteins were treated on ice for 20 min with 50 μg/ml of proteinase K. To inactivate the protease, 5 mM of PMSF was added to each reaction and the reaction was kept on ice for 10 min. Subsequently, the samples were centrifuged (20000 x g, 10 min, 4°C) and the pellets were mixed with 2x Sample buffer and boiled for 5 min at 95°C.

Import of TOM40 was monitored by resistance to alkaline extraction. To this aim, mitochondria were incubated for 30 min with 0.1 M Na_2_CO_3_ at 4°C. Then, samples were centrifuged (76000 x g, 30 min, 2°C), and the pellets were dissolved in 40 μl of 2x Sample buffer and heated at 95°C for 5 min before further analysis by SDS-PAGE. In some cases, the import of F1β and PINK1 was monitored by processing of the protein upon mitochondrial import. In these cases, at the end of the import reactions the mitochondrial pellets were directly resuspended in 2x Sample buffer.

Some import reactions were analysed by blue native (BN)-PAGE. In such cases, at the end of the import reactions the samples were centrifuged (13200 x g, 10 min, 4°C) and the pellets resuspended in digitonin-containing HMS-B buffer (3:1 (w/w) ratio of detergent to protein). Organelles were solubilized for 30 min at 4°C and non-solubilized material was removed via centrifugation (13000 x g, 10 min, 4°C). The supernatant was mixed with BN-PAGE sample buffer (5% w/v Coomassie blue G, 500 mM ∊-Amino-n-caproic acid, 100 mM Bis-Tris, pH 7.0) and samples were separated through a 4-16% polyacrylamide gradient gel. Radiolabeled proteins were detected by autoradiography.

### Expression and purification of recombinant proteins

Recombinant pSu9-DHFR-His was purified using a previously described protocol (Otera *et al.*, 2007). In brief, overnight culture of *Escherichia coli* BL21 cells expressing pSu9-DHFR-His was diluted in the morning to OD_600_ of 0.1 and once the culture reached an OD of 0.5, it was induced with 1 mM IPTG. The culture was grown for 3 h at 37°C and then cells were harvested by centrifugation. The pellet was resuspended in ice-cold buffer containing 50 mM Tris-HCl, pH 7.5, 150 mM NaCl as well as protease inhibitor cocktail (Sigma-Aldrich) and sonicated. The lysate was centrifuged to get rid of cell debris (15000 x g, 15 min, 4°C) and the clarified supernatant was applied to a nickel affinity resin column (Cube Biotech). The column was first washed with the buffer mentioned above supplemented with 10 mM imidazole. pSu9-DHFR-His was eluted in two steps using 40 mM and then 100 mM imidazole-containing buffer. To dilute out the imidazole and concentrate the sample, the eluted fractions were dialyzed against the same buffer lacking imidazole.

The purification of TcPINK1 was according to a protocol published by Rasool et al. (2018). The concentrated TcPINK1 was further purified by size exclusion chromatography (Superdex S200, GE Healthcare).

For purification of GST-fusion proteins, overnight cultures of *Escherichia coli* BL21 expressing GST alone, GST-Tom70, or GST-Tom20 were diluted in the morning to OD_600_ of 0.1 and, once the culture reached an OD of 0.5, it was induced with 1 mM IPTG. The cultures were grown for 4 h at 37°C and then the cells were harvested by centrifugation. The cells at the pellets were resuspended in ice-cold GST lysis buffer (20 mM HEPES, pH 7.25, 100 mM NaCl, 1.5 mM MgCl_2_, 2 mM PMSF, 1 mM DTT, 3 mM EDTA, protease inhibitor cocktail) and sonicated. Lysates were then centrifuged to remove cell debris (15 000 x g, 15 min, 4°C). Glutathione Sepharose 4B beads (GE Healthcare) were washed with GST basic buffer (20 mM HEPES, pH 7.25, 100 mM NaCl, 1.5 mM MgCl_2_) and incubated overnight at 4°C with the resulting supernatants. The beads with bound proteins were then transferred to purification columns. The columns were washed with GST basic buffer and subsequently, proteins were eluted with GST basic buffer containing 15 mM reduced L-glutathione. The fractions with the eluted proteins were then dialyzed to clear out L-glutathione and to concentrate the samples. Such purified proteins were then used in pull-down experiments. Tom70 used for peptide scan was further incubated overnight at 4°C with thrombin to cleave the GST tag. Afterwards, the GST moiety was rebound to the glutathione beads in the column and the flow-through containing Tom70 without the GST-tag was collected and used for further experiments.

### Immunodepletion of TOM complex from solubilized mitochondria

After import of PINK1 into depolarized mitochondria, the reactions were centrifuged (13200 x g, 10 min, 4°C) and the pellets were solubilized with digitonin as described above. Non-solubilized material was removed via centrifugation (13000 x g, 10 min, 4°C) and the resulting supernatant was incubated for 30 min at 4°C with antibodies against TOM22. Next, this mixture was incubated for another 30 min at 4°C with Protein A beads that were pre-washed with HMS-B buffer. The reaction mixtures were centrifuged (500 x g, 1 min, 4°C) and the supernatants were mixed with BN-PAGE sample buffer. Centrifuged protein A beads were washed three times with HMS-B buffer and proteins were eluted at 95°C for 10 min with 2x sample buffer.

### Peptide scan

Cellulose membranes harbouring 15-mer peptides covering the sequence of a.a. residues 1-140 of PINK1 were activated in methanol for 1 min and washed twice with sterile water for 1 min. Then, membranes were equilibrated at room temperature with binding buffer (50 mM Tris-HCl, 150 mM NaCl, 0.1% Triton X 100, pH 7.5) for 2 hrs. After blocking with 1 μM BSA for 1 h, membranes were incubated overnight with recombinant Tom70-cytosolic domain. Next, the membranes were washed three times with binding buffer and bound protein was visualized by immunodecoration using antibodies against Tom70.

### Pull-down of radiolabelled proteins with GST-fusion proteins

Radiolabeled proteins were synthesized as described above. Glutathione Sepharose 4B beads (GE Healthcare) were washed with GST basic buffer and incubated with purified GST-fusion proteins for 1 h at 4°C. Then, the reactions were centrifuged (500 x g, 1 min, 4°C) and supernatants discarded. Subsequently, the beads with bound recombinant proteins were blocked for 1 h at 4°C using reticulocyte lysate (Promega). Next, the beads were reisolated (500 x g, 1 min, 4°C) and incubated for 1 h at 4°C with radiolabeled proteins. ATP (0.2 mM) was added to the reactions every 30 min. At the end of the binding reactions, the samples were centrifuged again (500 x g, 1 min, 4°C) and supernatants discarded. The beads were then washed twice with GST basic buffer and pelleted again. Bound proteins were eluted at 95°C for 10 min with 2x sample buffer and analysed with SDS-PAGE followed by autoradiography.

## Acknowledgments

We thank E. Kracker for excellent technical assistance, N. Bartlick and A. Stasiak for help with some experiments, A.N. Bayne for help with the purification of TcPINK1, and V. Kozjak-Pavlovic for kindly providing cells. This research was supported by the German Research Foundation (DFG) through Research Training Group 2364 to K.M. and D.R. and through CRC 894 to M.J.

## Author contributions

K.K.M. designed and conducted experiments, M.J. prepared the peptide scan membranes,

S.R. J.F.T established a purification protocol for TcPINK1, D.R. designed experiments and analyzed data, K.K.M. and D.R. wrote the manuscript.

## Declaration of Interests

The authors declare no competing interests.

---

**Supplementary Figure 1.**
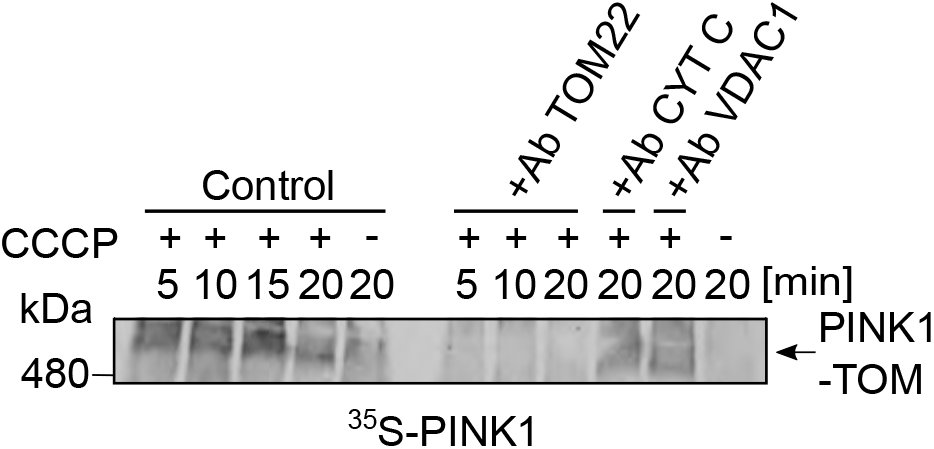
PINK1 associates specifically with the TOM complex upon depolarization of mitochondria. Radiolabeled PINK1 was incubated for the indicated time periods with isolated mitochondria, which in some cases were pretreated with CCCP. Organelles were then solubilized with digitonin and the lysate was incubated for 30 min at 4° C with antibodies against TOM22, Cytochrome C (CYT C), or VDAC1. Next, protein A beads were added for 30 minat 4°C and beads were pelleted. The supernatants were analyzed by BN-PAGE followed by autoradiography.

**Supplementary Figure 2.**
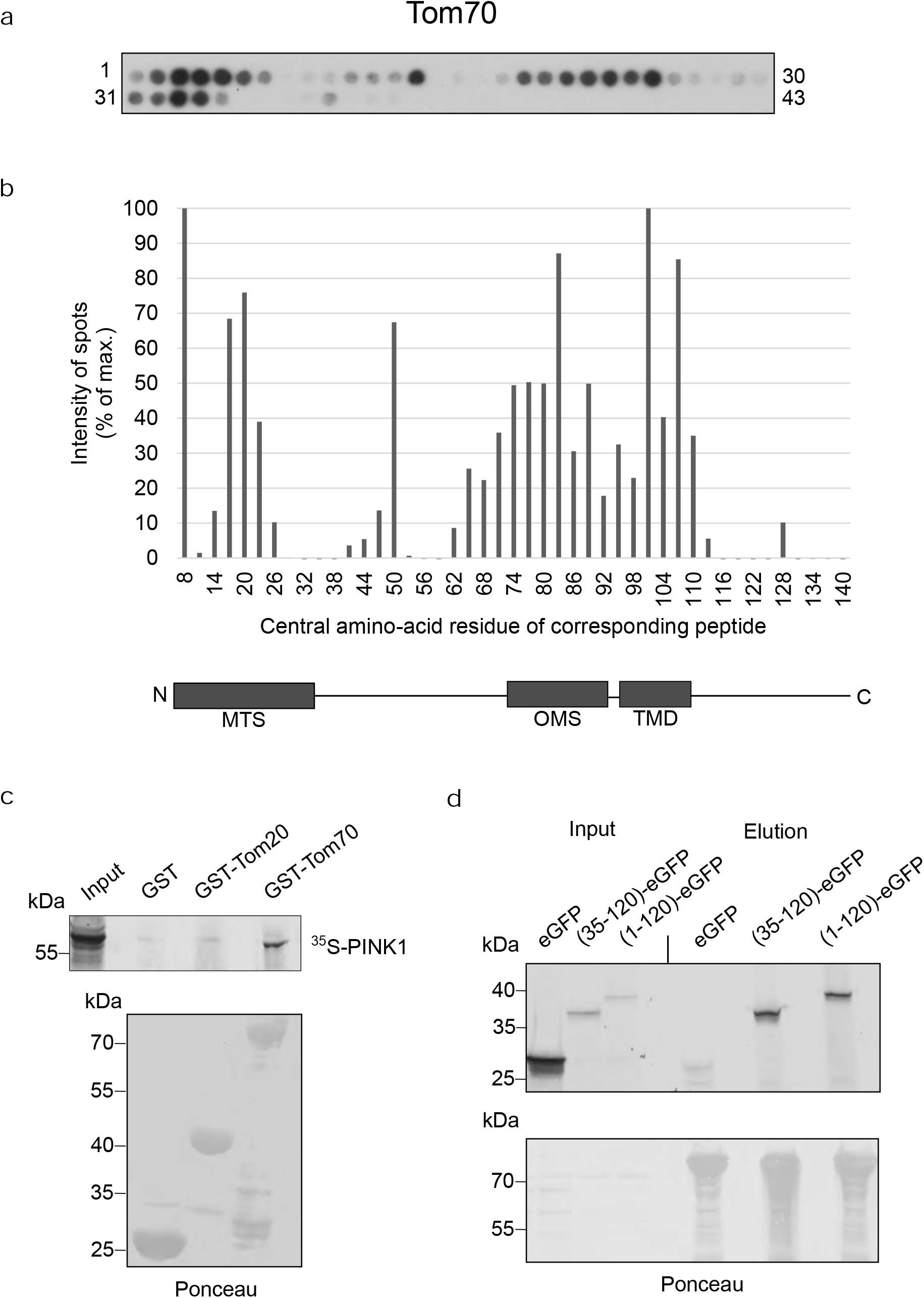
The cytosolic domain of Tom70 interacts directly with PINK1. **(a)** Nitrocellulose membrane containing 15-mer peptides covering amino acid residues 1-140 of PINK1 was incubated with recombinantly expressed cytosolic domain of yeast Tom70. After incubation, the interaction was visualized by immunodecoration using antibody against Tom70 (upper panel). The numbers flanking the panel reflect the serial numbers of the peptides. **(b)** The intensity of each dot was quantified and the average quantification of three independent experiments were plotted. The dot with the strongest intensity was set as 100%. The numbers on the X-axis reflect the central amino acid residue of each peptide. Lower panel: A schematic representation of the N-terminal region of PINK1 is shown. MTS, mitochondrial targeting signal; OMS, outer mitochondrial membrane localization signal; TMD, transmembrane domain. **(c)** GST, GST-Tom20, and GST-Tom70 were bound to glutathione beads and incubated with radiolabeled PINK1 for 1 h. Afterwards, proteins bound to the beads were eluted with 2x Sample buffer and analyzed by SDS-PAGE followed by blotting onto a nitrocellulose membrane. Radiolabeled proteins were detected by autoradiography whereas the recombinant GST-fusion proteins were visualized by Ponceau staining. **(d)** GST-Tom70 was bound to glutathione beads and incubated for 1 h with radiolabeled eGFP (as a control), PINK1(1-120)-eGFP, or PINK1(35-120)-eGFP for 1h. Afterwards, bound proteins were eluted with 2x Sample buffer and analyzed with SDS-PAGE followed by blotting onto a nitrocellulose membrane. Radiolabeled proteins were detected by autoradiography whereas recombinant GST-Tom70 was visualized by Ponceau staining.

